# WEADE: A Workflow for Enrichment Analysis and Data Exploration

**DOI:** 10.1101/252924

**Authors:** Nils Trost, Eugen Rempel, Olga Ermakova, Srividya Tamirisa, Letiția Pârcălăbescu, Michael Boutros, Jan U. Lohmann, Ingrid Lohmann

**Author notes:** Mailing address of corresponding author: Ingrid Lohmann, University of Heidelberg, Im Neuenheimer Feld 230, D-69120 Heidelberg, Germany, FX, EM.

## Abstract

Data analysis based on enrichment of Gene Ontology terms has become an important step in exploring large gene or protein expression datasets and several stand-alone or web tools exist for that purpose. However, a comprehensive and consistent analysis downstream of the enrichment calculation is missing so far. With WEADE we present a free web application that offers an integrated workflow for the exploration of genomic data combining enrichment analysis with a versatile set of tools to directly compare and intersect experiments or candidate gene lists of any size or origin including cross-species data. Lastly, WEADE supports the graphical representation of output data in the form of functional interaction networks based on prior knowledge, allowing users to go from plain expression data to functionally relevant candidate sub-lists in an interactive and consistent manner.

## INTRODUCTION

In the age of high-throughput sequencing technologies, there is increasing need for intuitive and consistent workflows that allow researchers making sense of the ever-growing data sets. The Gene Ontology (GO) annotation plays an important role in this, as it provides a controlled vocabulary of biological terms to which genes are annotated and that are familiar to wet-lab biologists [1,2]. For a descriptive analysis of gene sets at the functional annotation level, a variety of tools are available that determine the overrepresentation of GO terms, with the most common type of analysis being singular enrichment analysis (SEA) [3]. For this kind of analysis, the user must apply a threshold to the data (e.g. based on the p-value or fold change) to retrieve a set of candidate genes. This cut-off is highly arbitrary and always comes with the risk of losing potentially interesting candidates or including false positives. Other approaches perform gene set enrichment analysis (GSEA). In this case, no threshold has to be set, the complete data set is supplied and a statistic like the Kolmogorov-Smirnov test, or the Mann-Whitney U-test is used to calculate enrichment [4–7]. Both approaches have their up‐ and downsides, and the decision which to use should be on a case by case basis.

Since gene function is conserved across species boundaries, one conceptual advantage of GO analysis is that it can in principle be applied to compare datasets from diverse species. However, GO annotations for most species contain highly specific terms, which are not found in other annotations. Since most enrichment tools simply rely on the entire GO annotation set, cross species comparisons are virtually impossible due to non-matching annotations. There are several approaches to group terms and make results more amenable for comparative analyses: The simplest way is to restrict the analysis to high level GO terms, which represent very broad functional categories, such as “transcription” or “transport” found in all organisms. However, the information content of the terms on one specific level of the GO hierarchy is far from uniform across species [8]. Therefore, a better approach is to create a custom set of high-level terms, which are similar in information content, as measured by the number of annotated gene products, for diverse species. There are several such sets of terms, called GO slim ontologies that are available through the Gene Ontology Consortium. Seen from a biologist’s view, however, the terms in the generic GO slim still range from specific to general, including specific terms such as “transferase activity, transferring alkyl or aryl (other than methyl) groups” or very generic terms such as “reproduction”. Analysing enrichment results for such diverse terms across species boundaries is possible, but suffers from the inherent limitations in resolution and specificity of the resulting gene lists.

Apart from functional grouping by GO terms, sets of genes can be analysed by the identification of network structures through demonstrated genetic or protein-protein interactions. Tools that offer this functionality are for example GeneMANIA [9,10] and String [11,12], with BioGRID [13] being a database for gene-gene interactions. This information, taken together with the classification of genes into the Gene Ontology has the potential to represent a biological system comprehensively and intuitively.

Our WEADE tool offers a complete workflow of species independent GO enrichment calculation, data visualization and comparison, as well as functional network analysis. We describe the implementation of the tool, describe how WEADE can be used and compare it to a selection of currently available tools with similar functionality (GOrilla [14], g:GOST [15], GO_MWU [4–6], GSEA [7] and DAVID [3,16]).

## RESULTS AND DISCUSSION

Our aim was to develop a versatile and intuitive tool for genomic data analysis applicable to a wide variate of species and capable of cross-species analyses. We focused on the following issues: First, it should be possible to directly compare datasets derived from divergent experimental methods and/or organisms. Second, enrichment analysis should not be limited to simple candidate gene-lists, but should also allow the inclusion of all genes along with gene specific measurements, such as expression level or fold-change with the option to use these values for statistical testing within the tool. And third, the application should provide an intuitive workflow from uploading a dataset to exploring the results and testing hypotheses. This should include a user-friendly interface that can be used without prior knowledge and which would limit the necessity to change between tools. In the following paragraphs we will explain our approach and discuss in what ways our tool stands out from other gene set enrichment analysis tools currently available.

### Cross experiment – cross system analyses

To tackle the problem of comparability between species and experiments, we created a set of general functional annotation categories that are meaningful for any biological system. Each category consists of several GO terms that were handpicked to represent a certain biological function or property. These categories are not part of the GO tree itself and therefore encompass all three ontologies, biological processes, molecular functions and cellular components. Furthermore, the user may choose the ontologies that should be included in the analysis. The chosen categories allow for a comparison of different datasets irrespective of the model organism or the experimental method used to retrieve the data, as they are applicable to any of the supported biological systems. The categories are shown in Comparability between the categories can be judged by the number of annotated gene products across species in each category (table 1). Most of the assigned categories contain similar number of genes with the exception of the category ‘metabolism’, which stands out with over 200 million included genes. This is not surprising, since metabolism is not only a broad term, but also has diversified tremendously during evolution. In order to provide comparability between the categories, we normalized the enrichment results against the total number of annotated genes in each category (see Materials and Methods).

**Table 1.** High-level categories and numbers of annotated gene products per category. The numbers are taken from QuickGO and include annotations of all available species.

| Category | Number of annotations |
| --- | --- |
| **Cell cycle** | 1,999,523 |
| **Cytoskeleton, cell adhesion, and polarity** | 1,522,540 |
| **Differentiation** | 1,623,736 |
| **Growth** | 69,309 |
| **Immune** | 376,105 |
| **Metabolism** | 235,140,228 |
| **Reproduction** | 247,979 |
| **Signalling pathways** | 12,569,462 |
| **Stem cell** | 13,633 |
| **Stimulus** | 12,641,377 |
| **Trafficking and transport** | 33,264,077 |
| **Transcription factors and regulators** | 28,337,985 |
| **Translation** | 9,967,269 |

The first step of any WEADE analysis is based on the enrichment of these high-order categories rather than the enrichment of specific GO terms, making it possible to directly compare datasets from different species in a single workflow by simply selecting them from a dropdown menu. So far unsupported species can easily be analysed by uploading a custom background file that includes the annotations from gene identifiers to GO terms.

### Using input data

To calculate enrichment, we employ two different statistical tests, depending on the type of data submitted by the user. It is possible to use a defined list of candidate genes that were selected e.g. by thresholding for significantly over-expressed genes. In this case, we use the Fisher’s exact test. Alternatively, the user can provide the complete set of genes along with any data type that allows significance testing, e.g. expression value or fold change between experiments. We perform a Mann-Whitney U-test for this kind of data, avoiding the need to choose an arbitrary threshold for the data and potentially losing information.

The results of both types of analyses are represented in a plot and tabular form using intuitive color‐ coding for calculated p-value for the enrichment of the category. When calculating enrichment from a list of candidate genes using the Fisher test, the lengths of the bars represent the fraction of all annotated genes that appear in the sample and the category (Figure 1). In the case of calculating enrichment with the Mann-Whitney U-test (MWU), the lengths of the bars represent the delta rank, which indicates whether the category is predominantly in the lower or higher ranks (meaning down‐ or up-regulated in the example of using the fold change).

**Figure 1.**
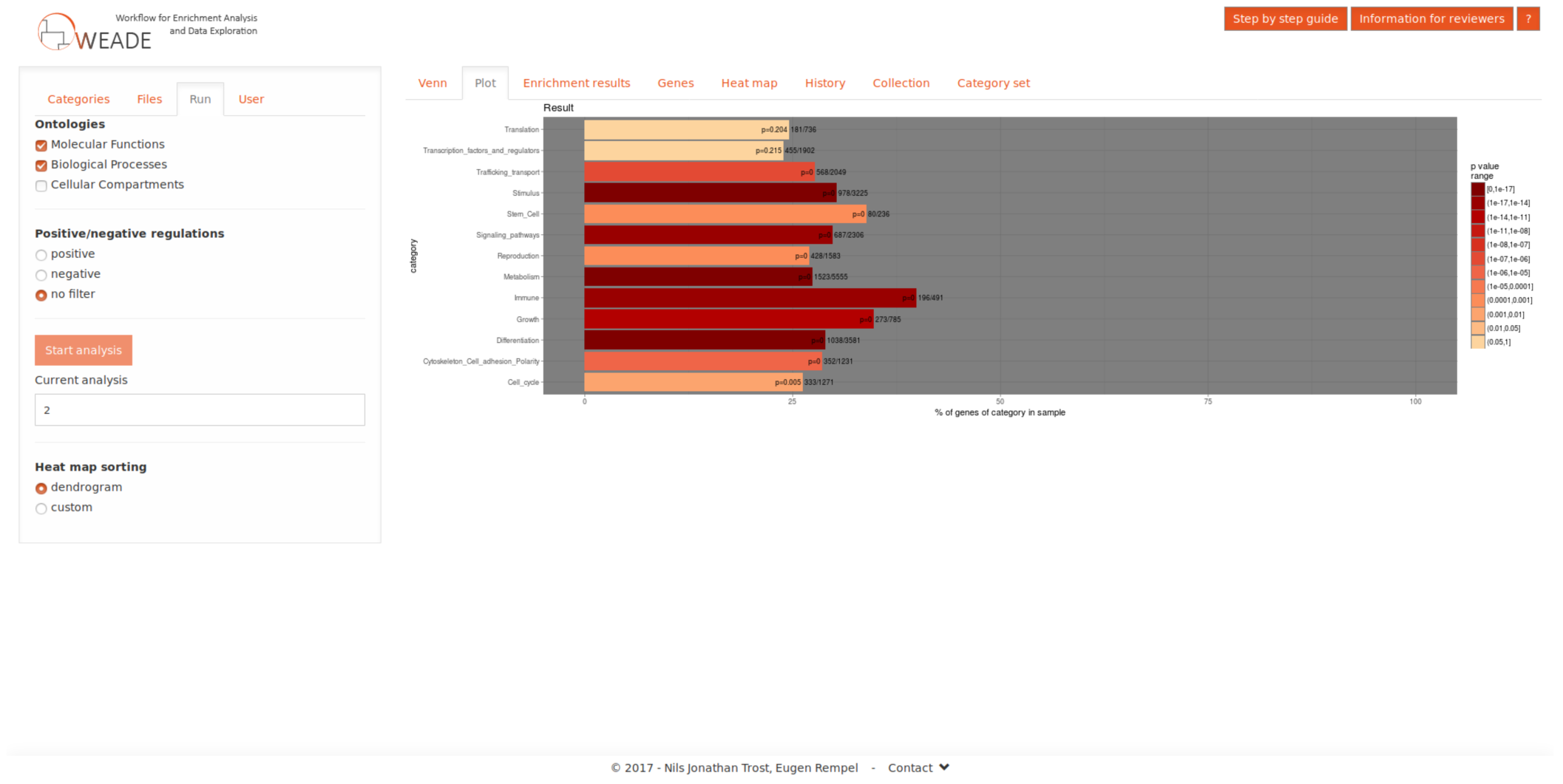
Screenshot showing the results section of the user interface after performing an enrichment analysis using a candidate gene set as input data. The colours of the bars represent the p-value of the enrichment, while the lengths of the bars represent the fraction of all annotated genes that appear in the sample and the category.

In addition to these results, we display a table with all genes in the data set and the categories to which they have been annotated. The genes are presented by their gene symbol for readability and include a link to the corresponding database, e.g. FlyBase for *Drosophila* data sets.

### Creating an integrated workflow

From this point on, the user can explore the data in a number of ways: One option is to run another parallel analysis on a related data set, and since an experiment often consists of several rounds using varying conditions users can upload up to four files. WEADE creates an intuitive output by generating a Venn diagram summarizing the overlaps of the genes in all uploaded sets, and any of the intersections can be chosen for further analysis. Importantly, the Venn diagram view also allows the iterative selection of a specific background, which results in enrichment calculation relative to another dataset rather than to all annotated genes, greatly facilitating comparative analyses of multiple experiments. The results of each round of analysis are stored during the session and can later be compared using an interactive heat map (Figure 2). The heat-map view can either display the p-value, or one of the measures specific to the statistical test, the normalized term-frequency in the case of the Fisher‘s exact test, or the delta-rank in the case of the MWU-test. Uploading new sets of data, or changing the background organism will not reset the heat map, but will create additional columns allowing iterative cross-species analyses.

**Figure 2.**
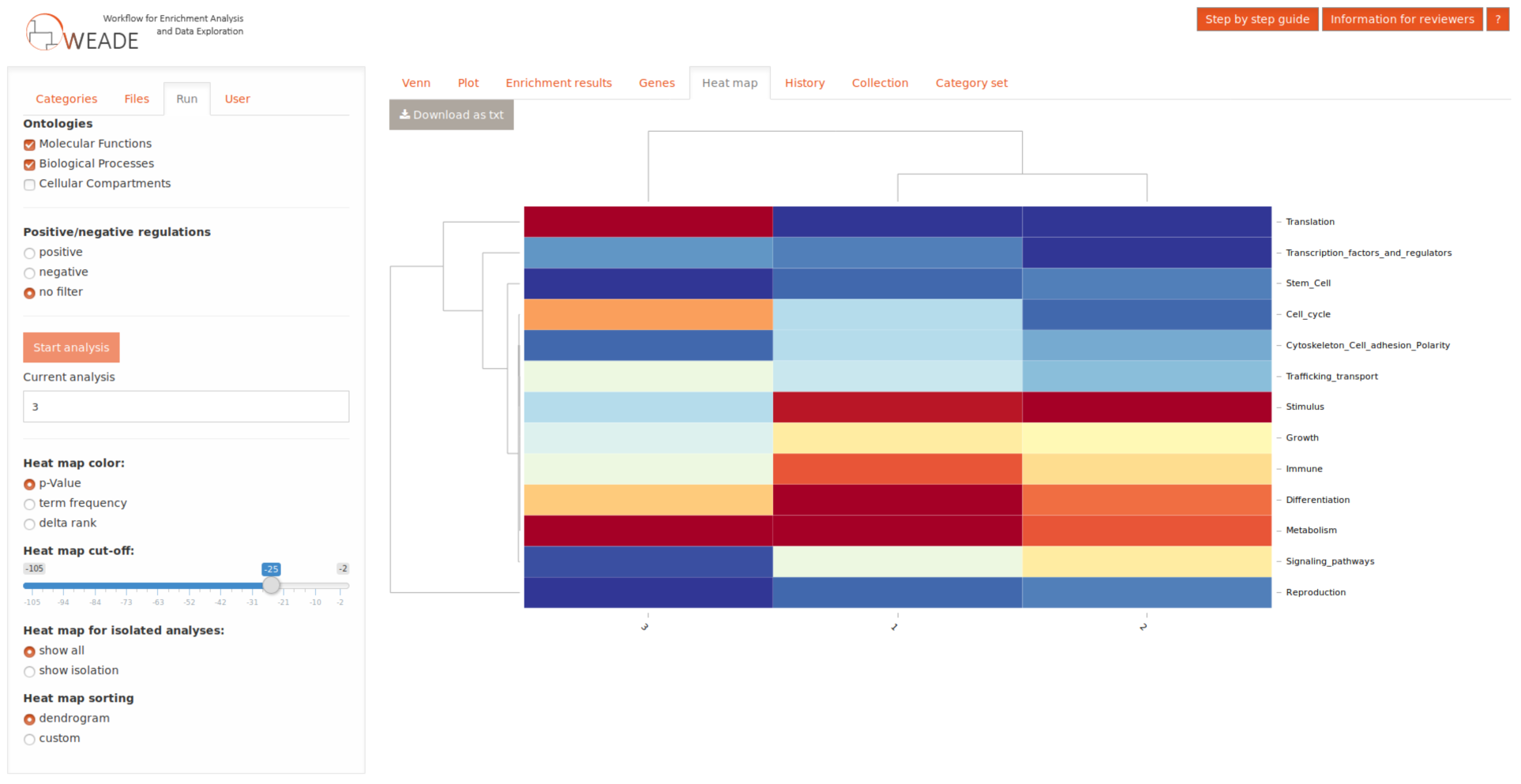
Screenshot of the interactive heat map that collects the results of previous analyses for easy comparison.

After getting an overview of the data with the broad categories, the user can then select a category and “zoom in” to see the individual GO terms that contributed to its enrichment. To this end, the enrichment analysis is performed again, but this time on the level of individual GO terms within the category. The results are represented in a sortable table, which includes links to QuickGO for convenient lookup of the GO term and a hierarchical network view for each term in the ontology tree. A table of contributing genes is also available for this analysis and all tables can be exported as tab separated text files for documentation or further analysis.

A third way to explore the data is by looking beyond functional annotations and including reported interactions between genes or proteins. To this end, users can select genes, either individually or from a GO term from a single species and visualize those in a functional interaction network (Figure 3). The genes or gene products are represented as nodes, while the edges correspond to the interactions, whose nature can be selected by the user. Genes from different analyses, or from different GO terms or selections can be displayed in a single network, with coloured nodes indicating the corresponding sets. We integrated an additional layer of information derived from Gene Ontology into the network by calculating the semantic distance between the GO terms of two genes to determine the length of the edge corresponding to the interaction between them. Here, we use a method loosely based on the method for detecting semantic similarity between GO terms described by Wang et al. [17]. However, we used a graph representation of the Gene Ontology and look for shortest paths, thereby we were able to limit the calculations making this type of analysis useful for a real-time interactive web application.

**Figure 3.**
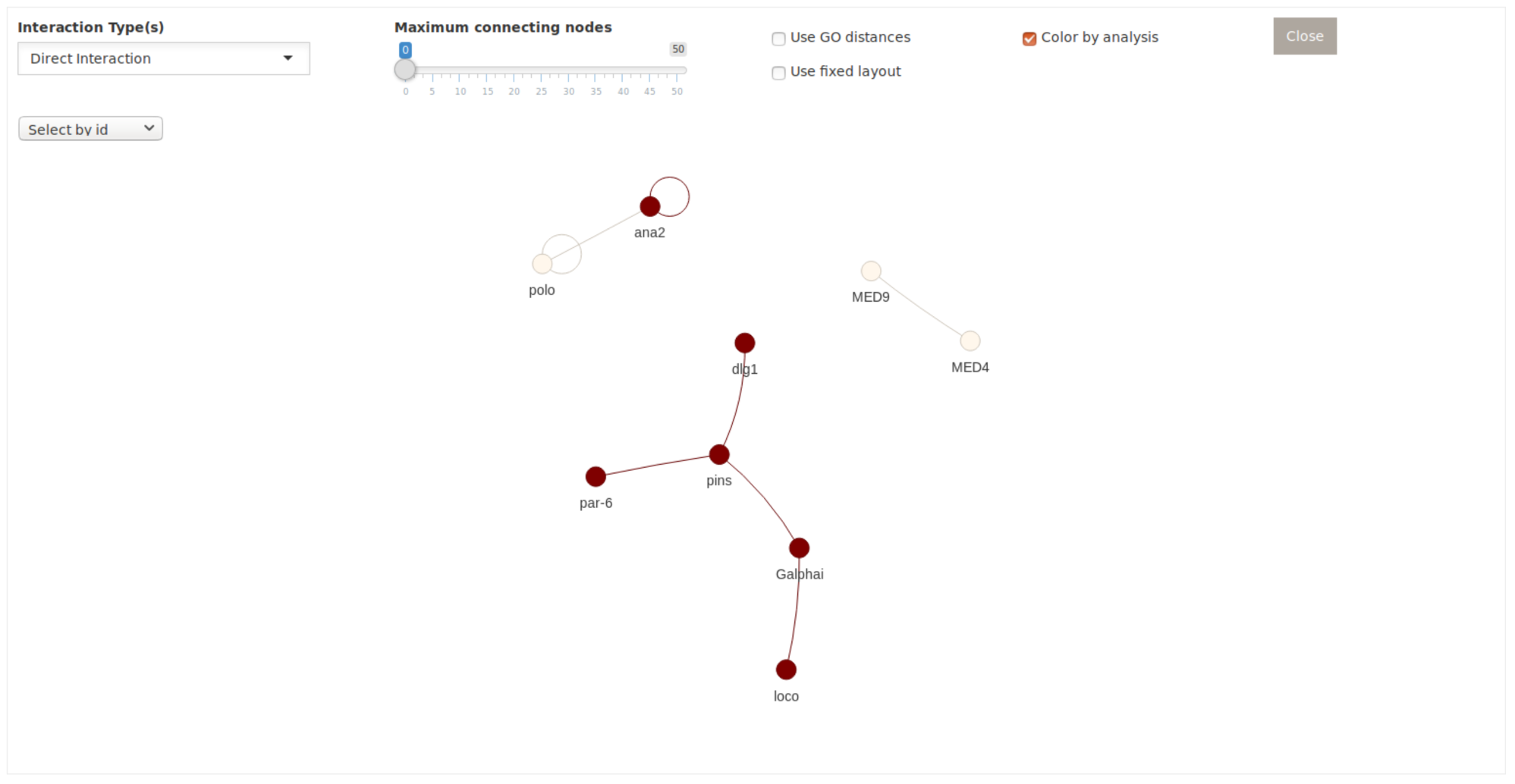
Screenshot of the user interface for the interaction network representation of genes. Differently coloured nodes come from different analyses.

Lastly, WEADE also leverages the BioGRID data to provide the option to identify receptor-ligand interactions between two datasets. This functionality is based on GO annotations to filtering for gene products acting as either receptors or ligands.

### Comparing WEADE to other tools

From the host of available tools for enrichment analysis, we have selected a few to compare their features to WEADE (see table 2 for a summary of the feature comparison). Some of the currently available tools are available as web applications, others need to be downloaded. Importantly, WEADE as a web application, works across all devices with internet access and is not limited by the computational resources or prior experience of the user. In the case of GSEA, the software needs to be downloaded and installed, GO_MWU comes in form of a set of scripts written in R and Perl and the user needs to be familiar with R to run the analysis.

**Table 2.** Feature comparison of tools with similar functionality.

| Tool | Enrichment Analysis |  |  |  | Interaction<br>Network | Availability |
| --- | --- | --- | --- | --- | --- | --- |
|  | SEA | GSEA | Graphic<br>Visualization | Result<br>Comparison |  |  |
| **WEADE** | Yes | Yes | Yes | Yes | Yes | Web Tool |
| **GOrilla** | Yes | No | Yes | No | No | Web Tool |
| **g:GOST** | Yes | No | No | No | No | Web Tool |
| **GO_MWU** | Yes | Yes | No | No | No | R script |
| **GSEA** | Yes | Yes | Yes | Yes | No | Software |
| **DAVID** | Yes | No | No | No | No | Web Tool |

Most currently available tools are limited to the analysis of candidate gene sets. This forces the user to either rely on an arbitrarily set threshold or test different thresholds, by manually sub setting the original data set multiple times. By additionally implementing the Mann-Whitney U-test, GO_MWU and WEADE allow the user to supply either a set of candidate genes or the complete data set with a continuous measure. GSEA has the same functionality in that regard, but using the Kolmogorov-Smirnov test instead of the MWU test. A key aspect of WEADE is the ability to quickly analyse multiple data sets and summarise the results in a heat map, thereby providing an on-site way to compare them. None of the web tools in this comparison include a similar feature, leaving only manual comparison by running the application several times and saving the results separately as an option for the user. GSEA is the only other tool to allow the comparison of multiple conditions inside the application, however requiring all data sets to use the same species annotation.

A feature which is uniquely present in WEADE, is the possibility to display genes in a functional interaction network. Displaying a set of interesting genes in an interaction network is a natural step during the analysis and would require the user to turn to other available tools, which include solely this functionality.

## CONCLUSION

Taken together, with WEADE we present a novel workflow that brings together frequently used tools for exploring data into a user-friendly, yet powerful web application. We plan on continuing the active development of the tool, expanding it in terms of supported species and additional features.

## MATERIAL AND METHODS

WEADE is an interactive web application built in R, HTML, JavaScript, and MySQL. The app is served with Shiny Server by RStudio. All data processing and analysis is done in R while the user experience is created using HTML pages and JavaScript. The underlying data structures are stored in a MySQL database.

### Creating GO Term Categories

The general set of categories was created by choosing a set of keywords that would represent biological areas in a consistent manner across species. These keywords were used to cluster GO terms from the generic GO slim subset into the general categories. The terms defining the categories were only allowed to be siblings in the GO graph; none of the terms are permitted to be offspring of other included terms. To avoid wrongly assigned terms, this automatic approach was followed by manually reviewing the categories, adding and removing terms where necessary. First and foremost, the categories should convey biological meaning.

### Enrichment Analysis

#### Analysis of candidate gene lists

To calculate the enrichment of GO term categories, a Fisher’s exact test is performed, where the genes with GO terms belonging to a category and included in the sample are compared against the total of genes in the background with GO terms of that category. The background may be a user selected set of genes or the complete annotation of genes of the selected species. The resulting p-values are adjusted for multiple hypothesis testing using the Benjamini-Hochberg method. In addition, the fraction of genes in the category and sample over background genes in the category is calculated.

When calculating the enrichment of single GO terms rather than of categories, the same method is applied, however, without p-value adjustment.

#### Analysis of gene lists with continuous measure

To calculate the enrichment of GO term categories or individual GO terms from a list of genes with a column of corresponding measurements, a Mann-Whitney U-test is performed. The p-value and the delta rank are reported. Like in the analysis of candidate gene lists, an adjustment to the p-value for multiple hypothesis testing is performed in the case of calculating the enrichment of categories.

#### Annotations

Annotations of gene identifiers to GO terms are retrieved from the respective biomarts. We use ensembl.org for animal annotations, plants.ensembl.org for plant annotations and fungi.ensembl.org for annotations of fungal species. Some select species annotations are available to date. These include *Drosophila melanogaster*, *Caenorhabditis elegans*, *Mus musculus*, *Danio rerio*, *Hydra vulgaris*, *Homo sapiens*, and *Arabidopsis thaliana*. A custom annotation file may also be used to expand the functionality of the tool to any species or for using an alternative gene identifier. This file can be uploaded using the advanced options in the file tab.

#### Gene interaction network

The current release of the BioGRID interaction repository is used to create a graph containing genes as nodes and their interaction as edges. The igraph R package is used to construct the graph and visNetwork for visualization. The user can select a set of genes to be presented in the network view. This selection is used to create a subset of the complete interaction graph. To visualize genes of the subgraph that are connected to one another not directly, but by one or more “connecting nodes”, a breadth-first search is conducted to find the shortest paths between the nodes in the complete graph. A user-defined number of connecting nodes can then be added to the subgraph. The user can choose what kind of interaction to display.

To add lengths to the edges, based on the relation of two connected genes in the Gene Ontology, a second graph is built containing the GO terms as nodes and their hierarchical relationships as edges. The length of the edge between Genes 1 and 2 is defined as the mean of all shortest undirected paths between each GO term of Gene 1 to all GO terms of Gene 2. A breadth-first approach is used to calculate shortest paths in the GO graph.

#### Pairwise interactions between datasets

The BioGRID data is searched for interactions, where interactor A appears in dataset 1 and interactor B in dataset 2. Both interacting genes must be annotated as either receptor (GO:0004872) or ligand (GO:0048018).

#### Documentation

An in-depth documentation of the tool and all its features is available from its homepage. Additionally, the tool includes a step-by-step guide to facilitate the first analysis.

#### Availability

The web application can be found at the following URL: http://weade.cos.uni-heidelberg.de

## Acknowledgement

We would like to thank the lab members of the I. Lohmann, J. Lohmann and M. Boutros labs for critically reading the manuscript

